# Longitudinal Analysis of Matched Patient Biospecimens Reveals Neural Reprogramming of Cancer-Associated Fibroblasts Following Chemotherapy in Pancreatic Cancer

**DOI:** 10.64898/2025.12.01.691614

**Authors:** Aylin Z. Henstridge, Nandini Arya, Ahmed M. Elhossiny, Padma Kadiyala, Georgina Branch, Vaibhav Sahai, Nicole Peterson, Jorge D. Machicado, Richard Kwon, Allison R. Schulman, Erik Wamsteker, George Philips, Stacy B. Menees, Jonathan Y. Xia, Tara L. Hogenson, Arvind Rao, Jiaqi Shi, Timothy L. Frankel, Filip Bednar, Marina Pasca di Magliano, Costas A. Lyssiotis, Mark J. Truty, Martin E. Fernandez-Zapico, Eileen S. Carpenter

## Abstract

Pancreatic ductal adenocarcinoma (PDAC), the most common subtype of pancreatic cancer, is a deadly disease with a complex tumor microenvironment (TME). How chemotherapy alters the TME, and whether these changes drive chemoresistance, is poorly understood. We examined matched pre- and post-treatment tissue specimens and found near-universal enrichment of axonal guidance genes in cancer-associated fibroblasts (CAFs) after treatment. These CAFs were enriched near sites of perineural invasion, coinciding with regions of increased tumor cell proliferation, and were enriched in tumor areas distant from nerves after chemotherapy. Metastatic recurrence lesions had the highest prevalence of these CAFs versus primary tumors and untreated metastasis. These CAFs showed elevated non-canonical Wnt mediators and axonal guidance genes, which complemented matching cognate binding partners in tumor epithelial cells, suggesting a role in tumor-stroma crosstalk. Our findings implicate fibroblast-derived axonal-guidance genes in promoting PDAC invasion and point to a promising target for this disease.

**Statement of Significance:** Therapeutic resistance remains a major challenge in pancreatic cancer. Our findings shed light on the changes in the complex tumor microenvironment in response to chemotherapy and identify fibroblasts high in axonal-guidance genes that may drive tumor progression and chemoresistance, thus uncovering a potential avenue for targeting pancreatic cancer.

## Introduction

Pancreatic ductal adenocarcinoma (PDAC) is a deadly disease predicted to be the second leading cause of cancer related death by 2040 (1,2). Diagnosis usually occurs at advanced stages of disease, making surgical resection available to only 15-20% of patients (3). The mainstays of therapy are cytotoxic combination treatments consisting of either FOLFIRINOX (5-Fluoroacil, irinotecan, leucovorin, and oxaliplatin) or gemcitabine/nab-paclitaxel. Unfortunately, most patients eventually progress due to chemoresistance, leading to poor outcomes (4). Even following surgical resection and chemotherapy, recurrence is common with a median overall survival of less than 2 years (1). Thus, there is an urgent need to elucidate mechanisms of therapeutic resistance to develop better strategies for treatment.

Although several datasets exist profiling treated and un-treated tumors (5–8), comprehensive studies exploring chemotherapy’s impact on the tumor microenvironment (TME) using longitudinal matched samples, thereby controlling for patient-specific variability, remain limited. This type of pancreatic tumor tissue sampling is resource-intensive, requiring a specialized endoscopic procedure. We have overcome this obstacle by obtaining treatment naïve endoscopic fine needle biopsy specimens at time of diagnostic biopsy and matched post-treatment specimens, at time of endoscopic fiducial placement or at surgical resection, to generate a unique longitudinal cohort of PDAC patients utilizing single-cell RNA sequencing (scRNA-seq). Notably, in contrast to many previous studies, processing was performed immediately on isolated live cells at time of tissue acquisition, allowing for robust profiling of both cytoplasmic and nuclear transcripts.

Here, utilizing these matched biospecimens, we define the TME landscape post-treatment. Transcriptomic analysis of the data revealed that changes in the immune microenvironment were heterogenous with minimal consistent alterations. In contrast, we observed consistent changes to cancer associated fibroblasts (CAFs) with an enrichment of neural axon guidance programs following treatment in primary tumors, and a further enrichment of this gene program in metastatic recurrence. These findings have important clinical implications given the hallmark of perineural invasion as a pathogenic process in PDAC (9–11), and the growing field of cancer neuro-pathobiology which investigates the role of peripheral nervous system (PNS) neurons as an active cellular compartment of the TME contributing to disease progression.

## Results

### scRNA-seq of matched pre- and post- chemotherapy tumors reveals diverse changes in the TME

To study the effects of chemotherapy in PDAC tumors, we collected longitudinal biospecimens from five patients by leveraging endoscopic fine-needle biopsies obtained during diagnostic workup. Following treatment with chemotherapy, we obtained biospecimens from either fine needle biopsies taken during endoscopic procedures performed for clinical care (fiducial placement) or procured tumor tissue from surgical resection (Figure 1A, B). The post-treatment samples for all patients were collected at least four-weeks after the conclusion of any chemotherapy treatment. Of note, one of the patients (1475) was additionally sampled at a third time point for tumor recurrence, taken 13 months after a total pancreatectomy (Figure 1A, B). The cohort was heterogeneous, both in terms of the treatment received (Figure 1B), and treatment response (Figure 1C and Supplemental Table 1A). Each sample was processed for scRNA-seq at time of collection.

**Figure 1.**
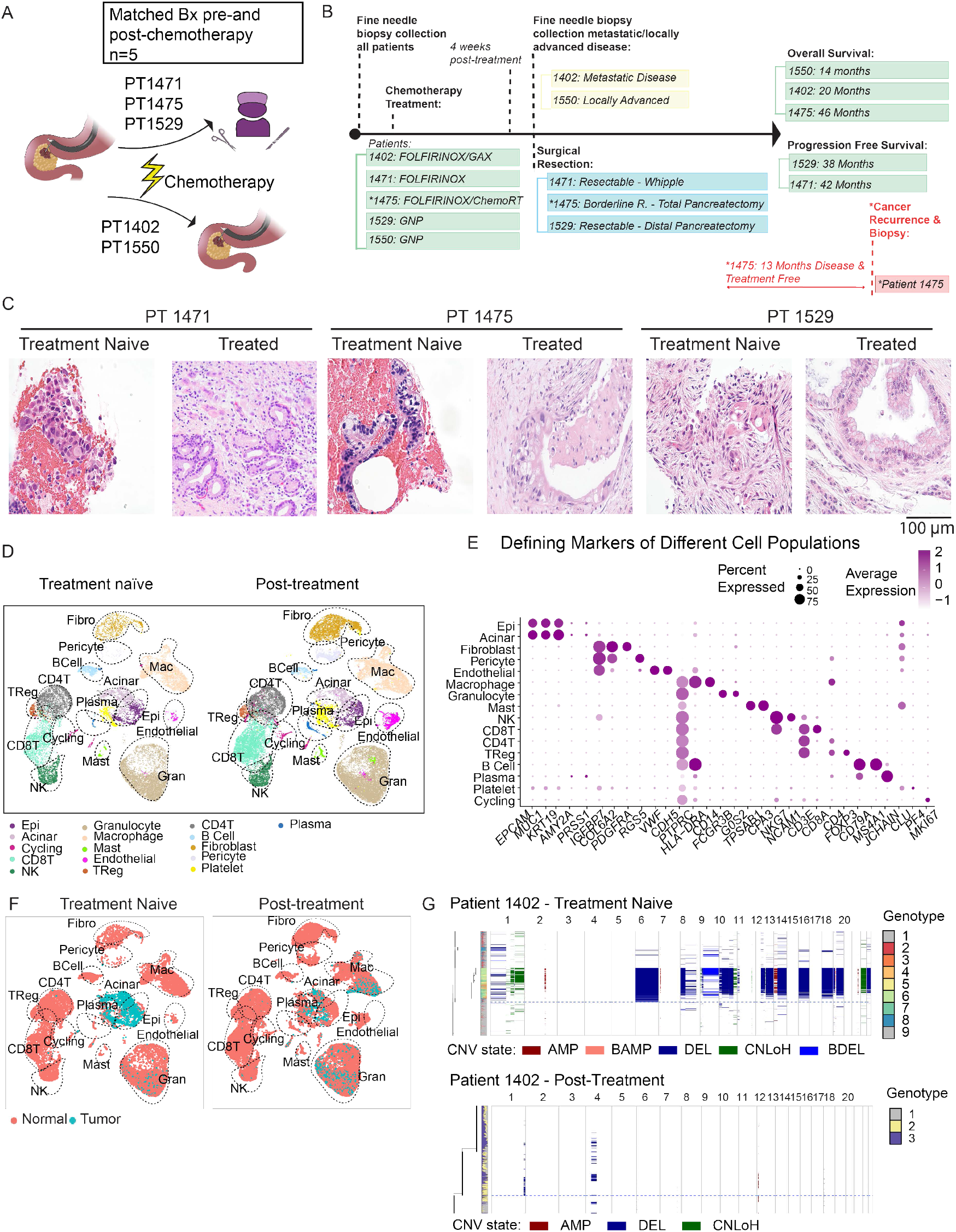
scRNA Sequencing of five matched patients pre- and post-chemotherapy reveals a diverse tumor microenvironment with transcriptomic variability. **A)** Schematic of samples from fine needle biopsies (FNB) or surgical resection. **B)** Timeline of patient sample collection, neoadjuvant chemotherapy treatments, and overall survival (for participants deceased) and progression-free survival (for patients currently being monitored) at time of data freeze. **C)** Hematoxylin and eosin (H&E) sections of representative biopsies (left) and resection samples (right) for each resectable patient in the cohort. **D)** UMAP of single-cell RNA sequencing (scRNA-seq) dataset generated from patient samples split by treatment naïve (left) and treated samples (right). **E)** Dot plot showing genes used to identify cell populations. **F)** UMAP reduction of scRNA-seq dataset with cells inferred to be ‘normal’ in red and ‘tumor’ in blue through inferred copy number variation analysis. **G)** Single-cell CNV landscape and reconstructed phylogeny of patient 1402 before (left) and after treatment (right). Legend indicates the identified variant as an amplification (AMP), balanced amplification (BAMP), deletion (DEL), balanced deletion (BDEL), or CNLoH (copy-neutral loss of heterozygosity).

To parse the cell-specific differences with treatment - epithelial, immune, and stromal cell clusters were manually identified using known gene signatures (Figure 1D, E). Pseudobulk analysis revealed broad transcriptomic differences between treatment naïve and post-treatment samples across all cell types, with no readily observable trends in cell type frequency across samples (Supplemental Figure 1A, B). We thus turned our focus to patient- and cell-type specific differences to parse out the effects of chemotherapy treatment.

Utilizing the *Numbat* (12) pipeline, which leverages allelic and gene expression frequencies to estimate copy number alteration, we identified cells within our population exhibiting a high likelihood of being malignant epithelial cells (Figure 1F). Upon observation of these proportions on a patient-specific level, it was determined that the cancer cell content of these biopsies post-treatment was too low to sufficiently analyze alterations in gene expression. This was likely due to the intended cytotoxic effects of chemotherapy (Figure 1G, Supplementary Figure 1C). The lack of malignant epithelial cells post-treatment was further corroborated by clinical pathologist examination of samples (Supplementary Table 1A, Figure 1C).

Following the absence of malignant cells to analyze post-treatment and given the hallmark immunosuppressive environment of PDAC (13,14), we next examined the immune landscape. NK and T cell populations were extracted and further classified based on previously published gene signatures (Figure 2A, B) (15). We queried known markers of immune activation and exhaustion within the T cell compartment for each patient and found that *TIGIT* expression in CD8+ T cells, specifically in exhausted CD8+ T cells (Supplementary Figure 2A), decreased following treatment in two patients (Figure 2C). Notably, these two patients had the best clinical outcomes in our cohort (Figure 1B, Supplemental Table 1A). One consistent finding was an increase in *CXCR4* expression in the CD8+ T cells of all patients and the CD4+ T cells of most patients post-chemotherapy (Figure 2C, D). Among the subpopulations of CD8+ and CD4+ T cells, *CXCR4* was enriched in all cytotoxic and exhausted CD8+ T cells, as well as regulatory T cells, with the steepest increase seen in cytotoxic CD8+ T cells (Supplementary Figure 2A). To understand the role of CXCR4 in the context of treatment, we used the CellChat V2 pipeline (16) to look at inferred ligand-receptor interaction strength and specifically queried chemokine signaling in the immune-stromal compartment. We identified a post-treatment increase in *CXCL12-CXCR4* interactions between CAFs and both CD8+ and CD4+ T cells (Figure 2E, F). Of note, *CXCR4* was not found to be appreciably expressed in CAFs in our cohort (Figure 2E). The *CXCL12-CXCR4* axis has been shown to play an important role in suppressing T cell responses through impairing immune cell directed migration (14,17–19). Our findings suggest that this crosstalk may be enhanced after chemotherapy treatment, thereby serving as a mechanism for chemoresistance.

**Figure 2.**
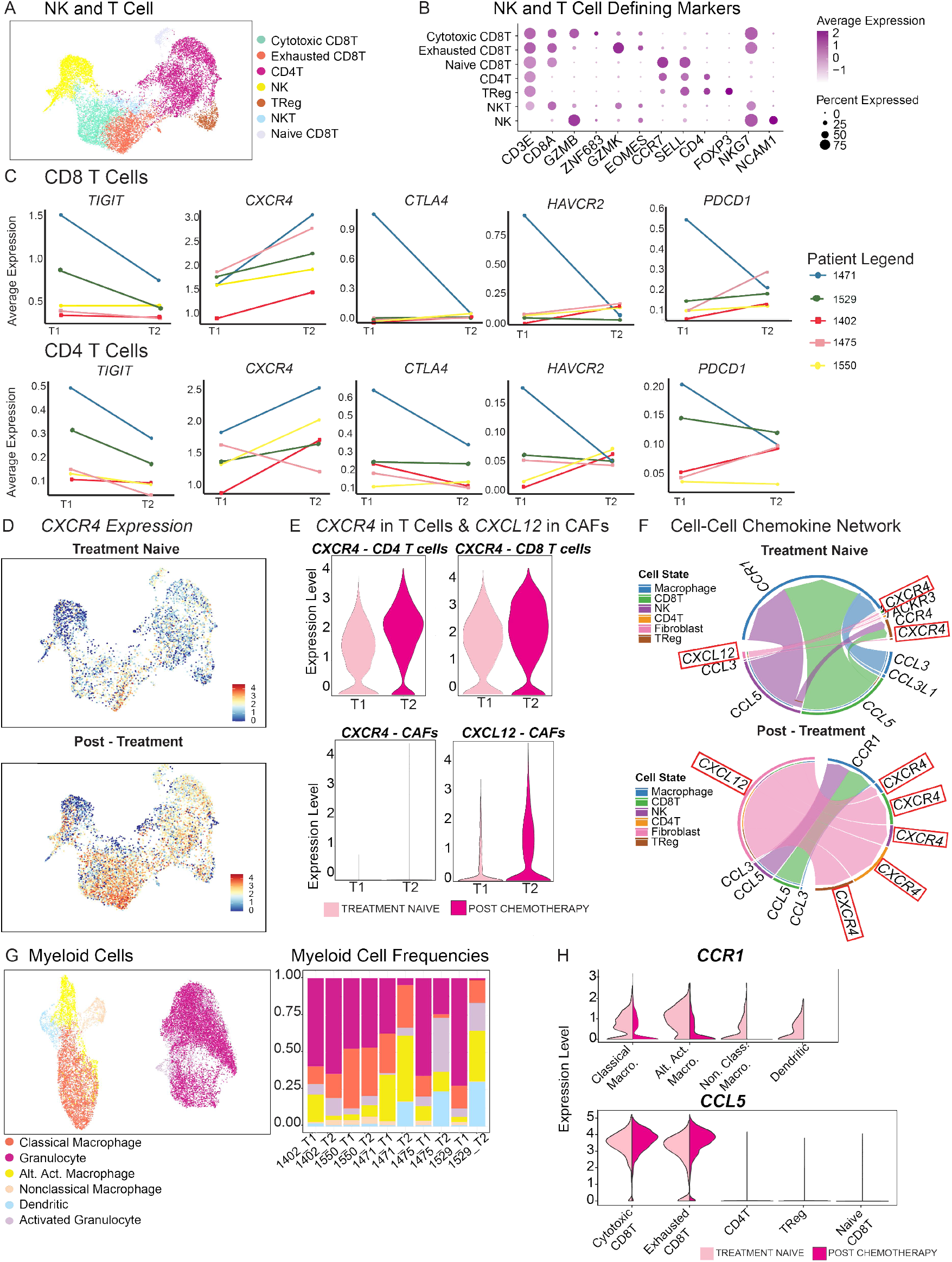
CXCR4 is unanimously enriched CD8+ T cells post-treatment, with increased interactions between T cells and cancer associated fibroblasts. **A)** UMAP reduction of extracted T cells and NK cells. **B)** Dot plot showing genes used to identify T cell and NK sub-populations. **C)** Before-and-after plots of immune-cell markers in each patient (CD8+ T cells, top and CD4+ T cells, bottom). **D)** Feature plot showing gene expression of *CXCR4* in all T cells and NK cells in UMAP reduction of average gene expression scaled from 0 to 4 split by treatment naïve (top) and treated (bottom) t cells of all patients. **E)** Violin plots showing average *CXCR4* gene expression in CD4 and CD8 T cells and average *CXCR4* and *CXCL12* gene expression in cancer associated fibroblasts split by treatment status. Adjusted *P* value for significantly differentially expressed markers: *CXCR4 in T cells < 2*.*2e-16 and CXCL12 in CAFs < 2*.*2e-16*. **F)** Circle plots showing inferred ligand-receptor pairs between macrophages, T cells, NK cells, and fibroblasts in treatment naïve (top) and treated (bottom) samples from all patients. **G)** UMAP of identified myeloid cell populations and cell frequency plot identifying fraction of cell subtypes in each population split across patient samples in treatment naïve and treated populations. (Alt. Act. Macrophages are alternatively activated macrophages). **H)** Violin plots showing *CCR1* expression in all myeloid cells split by time point (top) and *CCL5* gene expression in all T cells split by time point (bottom).

Turning our attention to the myeloid compartment, which contained macrophages, granulocytes, and dendritic cells, we found an increase in alternatively activated macrophages following chemotherapy (Figure 2G, Supplementary Figure 2B). We previously reported this population, marked by *C1QA, APOE*, and *SPP1*, to be important for establishing the premetastatic niche (Supplementary Figure 2B) (20). Interestingly, highly enriched in this population pre-treatment, we saw a decrease in *CCR1* expression, and a corresponding decrease in *CCR1-CCL5* signaling between macrophages and T cells/NK cells (Figure 2F, H). CCL5 signaling to macrophages has been reported to polarize them into a tumor-promoting and immunosuppressive subtype (21,22). Altogether, our observed alterations in the immune microenvironment show that chemotherapy impacts specific cell compartments in diverse ways - potentially driving context-dependent immunosuppressive and immune-activating effects.

### Chemotherapy induces the enrichment of CAFs with an ‘axonal guidance’ gene signature

The increase in CAF-derived *CXCL12* post-therapy prompted us to further examine the CAF population in our dataset. This yielded three distinct clusters of CAFs - which we termed Fibro1, Fibro2 and Fibro3 (Figure 3A, B). These populations were marked by varying expression levels of canonical CAF genes (*ACTA2, PDGFR*α, *TAGLN, FAP*), as well as *CXCL12* identified predominately in Fibro1 and Fibro2 (Figure 3C, D). Interestingly, the top expressed genes in Fibro3 were genes associated with axonal guidance and neurite outgrowth (*NRP1, EPHA3, NEGR1, ROBO1, SLIT2, NFIA, SEMA3A*) (Figure 3C, D). These genes were particularly of interest due to their roles in axon guidance, as well as past associations with neural invasion and treatment resistance in cancer (23–25). We thus developed a 7-gene signature consisting of these axonal guidance, henceforth referred to as the neural-CAF signature (Figure 3C-E). In order to better understand the relationship of these genes to previously reported CAF signatures, we applied published myofibroblast (myCAF), inflammatory fibroblast (iCAF), and antigen-presenting fibroblast (apCAF) gene signatures to our dataset (Figure 3E) (26).

**Figure 3.**
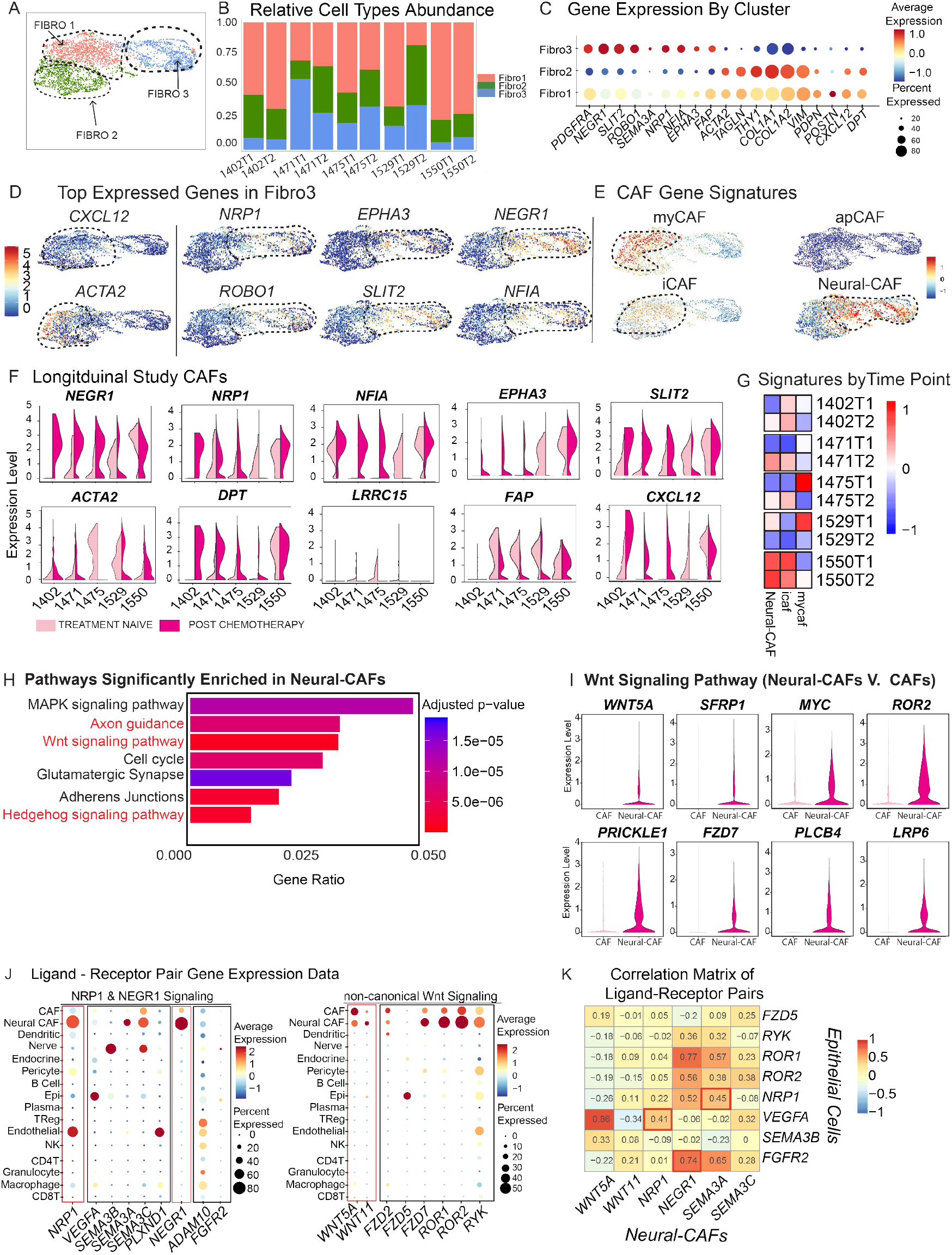
An ‘axonal guidance’ gene signature is enriched in a sub-population of CAFs following treatment. **A)** UMAP reduction of extracted fibroblasts. **B)** Histogram showing cell frequencies for Fibro1, Fibro2, and Fibro3, split by treatment status (T1 – treatment naïve and T2 – treated). **C)** Dot plot showing top expressed genes in Fibro1, Fibro2, and Fibro3 populations. **D)** Feature plot of fibroblast UMAP reduction showing average gene expression of *CXCL12, ACTA2* (left) and top expressed genes in Fibro3 (right). **E)** Feature plot genes representative of AUCell gene signature scoring of myofibroblasts (top left), inflammatory fibroblasts (bottom left), and antigen presenting fibroblasts (top right), and UMAP reduction of fibroblasts with AUCell scoring of our neural-CAF gene signature (*NEGR1, NRP1, NFIA, ROBO1, SLIT2, SEMA3A, NFIA, EPHA3*) (bottom right). **F)** Violin plots of average expression of neural-CAF and common fibroblast genes split by patient and colored by treatment status. **G)** Heatmap of signature scores of neural-CAF gene signature, myofibroblastic gene signature, and inflammatory fibroblast gene signature split by patient ID and treatment status (T1 – treatment naïve and T2 – treated) (right). **H)** Bar plot showing enriched pathways from the KEGG database comparing neural-CAF to CAFs. X-axis identifies gene ratio, and bar color indicates p-value. Pathways of interest are marked in red. **I)** Violin plot of top differentially expressed genes for Wnt signaling in neural-CAF and CAFs. Adjusted *P* Value for significant genes: *WNT5A: 0*.*21, SFRP1: 2*.*8e-10, MYC: 2*.*9e-12, ROR2 <2*.*2e-16, PRICKLE1 <2*.*2e-16, FZD7 <2*.*2e-16, PLCB4 <2*.*2e-16, LRP6 <2*.*2e-16*. **J)** Dot plot of ligand-receptor pairs involved in NRP1, NEGR1, and non-canonical Wnt signaling in all cell subtypes - CAF (cancer associated fibroblast), Endo. (Endothelial cells), Peri. (Pericyte), Gran. (granulocyte), Epi. (epithelial) **K)** Correlation matrix between ligand-receptor pairs expressed in either neural-CAFs or Epithelial cells. X-axis contains genes expressed in neural-CAFs and y-axis contains genes expressed in epithelial cells. Each cell displays the degree of correlation between genes expressed in each population. Correlations of interest are highlighted with red boxes.

The myCAF and iCAF signature mapped to partially overlapping regions in Fibro1 and Fibro2, consistent with prior reports that fibroblasts can express both characteristics simultaneously (27,28). apCAF genes were identified in a few individual cells but did not identify a separate cluster. While the Fibro3 cluster was most enriched for the neural-CAF signature, there was overlap between cells positive for the neural-CAF and iCAF signature, potentially suggesting an intermediate state (Figure 3E). To further investigate changes at a patient-specific level, we observed that many components of the neural-CAF signature increased in expression post-treatment (Figure 3F). Additionally, we saw the opposite trend with *ACTA2*, suggesting a decrease in myCAFs after chemotherapy treatment, as well as shifts in fibroblast gene markers (Figure 3F). Specifically, *FAP* and *LRRC15* (29,30), both associated with immunosuppression in the TME, were decreased following treatment, while *DPT*, a pan-fibroblast marker, found to be expressed variably across fibroblast populations (31), was increased (Figure 3F). We further compared the neural-CAF, myCAF and iCAF gene signatures patient-by-patient across treatment groups and found a dominant trend of increased iCAF and neural-CAF expression with decreased myCAF expression (Figure 3G). Patient 1529 was consistently an outlier to this trend, but it is worth noting that this patient had displayed the best response to treatment out of the cohort (Figure 1B, Supplemental Table 1A).

Based on these findings, we interrogated the neural-CAF signature in our previously published dataset of untreated PDAC tumor samples (14). Even with the inclusion of more treatment-naïve PDAC CAFs, enrichment of the neural-CAF signature remained highest in our post-treatment samples (Supplementary Figure 2C). Notably, we mapped the neural-CAFs in this larger dataset and identified that they not only were present but clustered separately from the myCAF population (Supplementary Figure 2D). In this larger cohort, we queried enriched KEGG pathways and differentially expressed genes between neural-CAFs and the remaining CAF subtypes. In addition to enriched axonal guidance, neural-CAFs were enriched for Wnt and hedgehog signaling (Figure 3H). We also observed that they were specifically enriched for genes such as non-canonical Wnt Ligand *Wnt5A*, receptors *ROR2* and *FZD7*, and downstream targets such as *Myc* (Figure 3I). Of note, neuronal signaling involving NRP1 has been previously associated with Wnt and Hedgehog signaling targets (32). We queried the expression of canonical ligand-receptor pairs enriched in individual cell populations from axonal guidance and Wnt signaling pathways (Figure 3J). The cognate ligand-receptors for each of these pathways were most strongly expressed in neural-CAFs and epithelial cells, as well as endothelial cells (Figure 3J). We further observed that the gene expression of a subset of these cognate ligand-receptor pairs (*SEMA3A-NRP1, NRP1-VEGFA, NEGR1-FGFR2)* were correlated between neural-CAFs and epithelial cells (Figure 3K). Altogether, we observed that chemotherapy exerts diverse changes on the immune microenvironment in PDAC while consistently promoting the upregulation of neuronal axon guidance genes in CAFs, which may contribute to tumor survival through Wnt signaling.

Observing that the neural-CAF signature is present among CAFs in naïve tumors and enriched post-treatment, we further queried the expression of these genes in the absence of disease pathology using our published scRNA-seq dataset of healthy pancreata (31). Upon comparing healthy pancreatic fibroblasts and PDAC associated fibroblasts, we observed that a subset of these neural genes (*NEGR1, NRP1, NFIA*, and *SLIT2)* were present in the fibroblasts of healthy tissue (Figure 4A). *EPHA3*, however, was specific to CAFs. As expected, and previously validated by others (31), *FAP* and *ACTA2* were higher in CAFs, while *PDGFR*α, *PDGFR*β, *DPT*, and *CXCL12* were expressed in both populations (Figure 4A). For further validation, we performed co-immunofluorescence staining of donor pancreas tissue for PDGFRα, NRP1, and NEGR1 (Figure 4B, C and Supplementary Figure 3A-C). We observed the presence of these neural genes in fibroblasts in healthy donor pancreatic tissue, specifically near pancreatic ducts (Supplementary Figure 3B, C), pancreatic intraepithelial neoplasia (PanIN) (Figure 4B, Supplementary Figure 3A), and the epineurium of nerves within the healthy pancreas (Figure 4C). These findings suggest that while elements of the neural-CAF signature are present in healthy pancreatic fibroblasts, elements of this transcriptional program are present only in the TME and are further enriched following chemotherapy.

**Figure 4.**
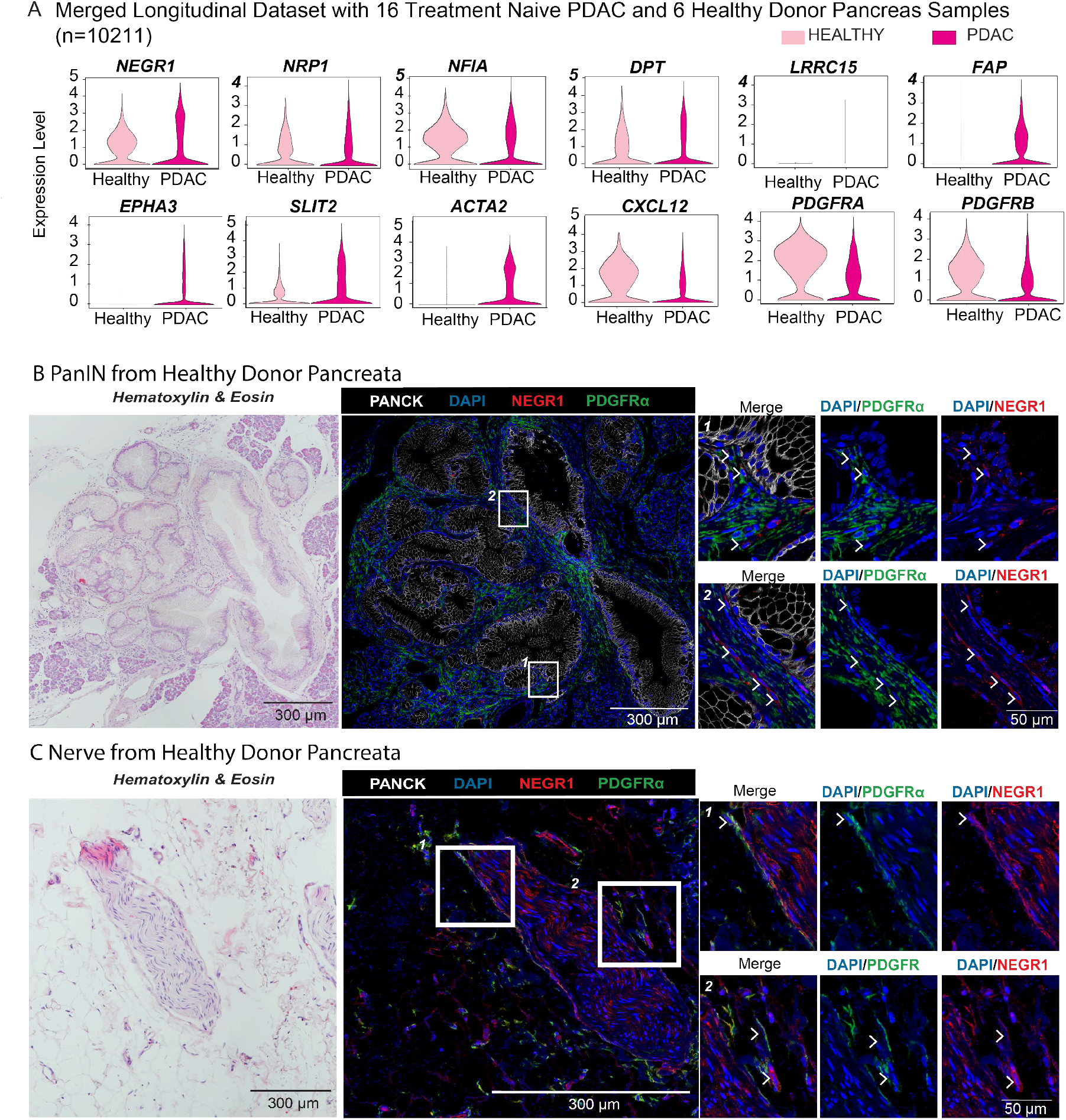
The neural-CAF signature is partially recapitulated in the absence of malignancy. **A)** Violin plots of CAF markers and neural-CAF genes comparing healthy fibroblasts from healthy donor pancreata and PDAC samples. Adjusted *P* value for genes: *NRP1 0*.*067, NEGR1 5*.*5e-05, DPT 0*.*00012, NFIA <2*.*2e-16, FAP <2*.*2e-16, EPHA3 <2*.*2e-16, SLIT2 <2*.*2e-16, ACTA2 <2*.*2e-16, CXCL12 <2*.*2e-16, PDGFRA <2*.*2e-16, PDGFRB <2*.*2e-16*. **B)** Hematoxylin and eosin (HE) staining of PanIN identified in healthy pancreas (left) and stained for DAPI (blue), PDGFRα (green), PANCK (white), and NEGR1 (red). White arrows denote NEGR1+ fibroblasts at higher magnification (right). **C)** HE of tissue resident nerve in healthy pancreas (left) stained for DAPI (blue), PDGFRα (green), PANCK (white), and NEGR1 (red). White arrows denote NEGR1+ fibroblasts at higher magnification (right).

### Neural-CAFs preferentially localize near sites of perineural invasion, while their abundance is specifically elevated in nerve-distant tumor regions following chemotherapy

As scRNA-seq does not preserve spatial information, we performed co-immunofluorescence staining on treatment naïve and treated patient PDAC tissue to identify the spatial distribution of neural-CAFs. Under guidance of a trained pathologist, we identified areas of perineural invasion, tumor cells (distant to nerves), and nerves (without adjacent tumor cells in proximity) within each patient sample (Supplementary figure 4A, B).

Using αSMA as a fibroblast marker, we quantified the amount of NEGR1+ CAFs present in the surrounding stroma of each of these structures. The percent of NEGR1+ CAFs were quantified utilizing radial bins 0-300 μm away from nerve or tumor ducts in 50 μm intervals (Supplementary Figure 4C). We observed that NEGR1+ CAFs were more abundant near nerves, particularly in areas of identified perineural invasion (Figure 5A-C).

**Figure 5.**
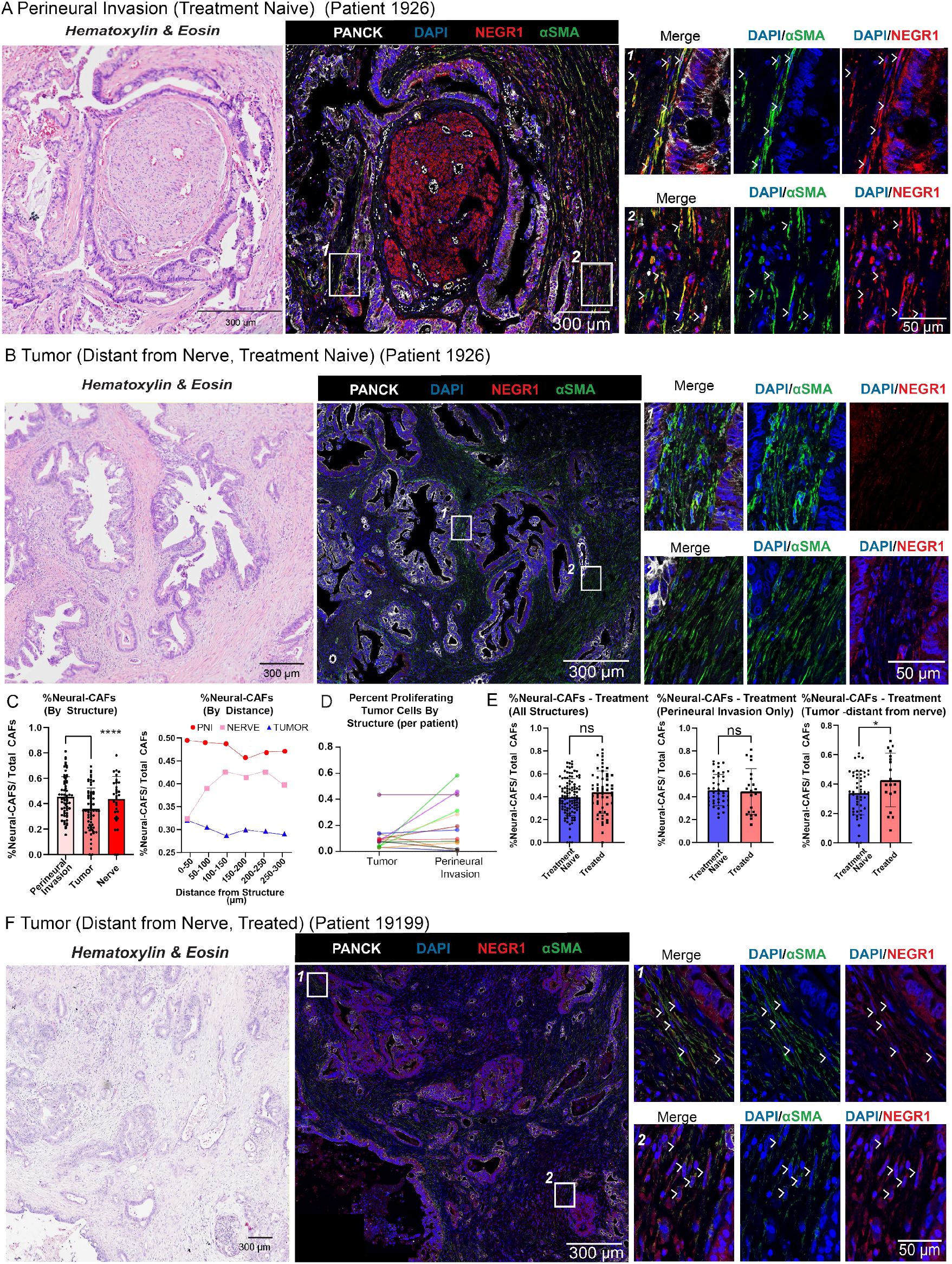
Neural-CAFs are enriched near regions of perineural invasion in both treatment-naïve and treated PDAC, whereas their presence in tumor areas distant from nerves is specifically elevated following chemotherapy. **A)** Hematoxylin and eosin (HE) staining of treatment naïve patient tissue (patient 1926) (left), and immunofluorescence staining of serial section from same patient tissue with DAPI, aSMA (green), PANCK (white), and NEGR1 (red) in an area marked as perineural invasion (PNI) (middle). White arrows denote NEGR1+αSMA+ CAFs at higher magnification (right). **B)** HE staining of treatment naïve patient tissue (patient 1926) (left), and immunofluorescence staining of serial section from patient with DAPI, aSMA (green), PANCK (white), and NEGR1 (red) in tumor tissue non-adjacent to PNI. White arrows denote NEGR1+αSMA+ CAFs at higher magnification (right). **C)** Bar plot of percentage of neural-CAFs in regions of PNI, tumor (distant to nerve), and nerve alone from each patient (N=17) with N=10 random regions of interest chosen for each patient. Spatial distribution plot showing percentage of neural-CAFs out of total CAFs in each structure at distance bin of 50 μm intervals (right). Adjusted P value for %neural-CAFs in areas of PNI versus areas of tumor cells (distant from nerve) *<0*.*0001*. **D)** Quantification of average %KI67+PANCK+/Total PANCK+ cells in areas of PNI and areas of tumor alone, where each line represents one patient (paired t-test *P=0*.*07*). **E)** Bar plot of percentage of neural-CAFs in treatment naïve patients (N=11) and treated patients (N=6) with all structures included (left), only areas of PNI (middle), and only areas of tumor (distant to nerve) (right). Adjusted *P* values for treatment naïve versus treated patients: in tumor cells (distant to nerve) *0*.*0416*, all structures combined *0*.*1258*, and PNI only *0*.*7671*. **F)** HE staining of treated patient tissue (patient 191999) (left), and immunofluorescence staining of serial section from same patient tissue with DAPI (blue), αSMA (green), PANCK (white), and NEGR1 (red) in an area marked as tumor (distant to nerve) (middle). White arrows denote NEGR1+αSMA+ CAFs at higher magnification (right).

Interestingly, we further observed that areas of perineural invasion in patient tissues trended towards having higher levels of proliferating tumor cells compared to tumor areas non-adjacent to perineural invasion, regardless of treatment status (Figure 5D, Supplementary Figure 4D). While the percent NEGR1+ CAFs was high in areas of perineural invasion regardless of treatment status, we did see a difference in abundance of NEGR1+ CAFs with treatment status in areas of tumor glands distant to nerves, with significant enrichment post-treatment (Figure 5E, F). We also validated these findings with a second neural-CAF marker, NRP1, and found similar results (Supplementary Figure 5A-C). Furthermore, analysis of the spatial distribution relative to nerve structures revealed that abundance of neural-CAFs was highest within 200 μm from tumor-invaded nerves while nerves in the absence of any perineural invasion showed a reciprocal distribution (Figure 5C).

This was likely in part due to nerves in isolation having less stromal cells surrounding them compared to areas with tumor cells present. Altogether, we found that neural-CAFs are enriched in areas of perineural invasion, irrespective of treatment status. In contrast, in areas of tumor distant to nerves, neural-CAFs are enriched in post-treatment samples compared to treatment naive tumors. These findings indicate that the acquisition of the neural-CAF signature in fibroblasts may result from tumor-nerve interactions, or alternatively, may be promoted by the selective pressures of chemotherapy.

Given the trends of neural-CAF enrichment in tumors with treatment, we sought to query the potential effect of these CAFs on the tumor cells themselves. Of unique interest to us was patient 1475, from whom we obtained a third point for analysis in the setting of cancer recurrence. Patient 1475 received neoadjuvant treatment and chemoradiation followed by a total pancreatectomy. Surgical pathology showed a near complete response to treatment, reflected in our single cell sequencing sample which captured no tumor cells (Figure 6A, Supplementary Table 1A, Supplementary Figure 1C). However, 13 months post resection, this patient developed a para-aortic mass which was found to be a recurrence of their disease when biopsied (Figure 6A). We observed that the inferred copy number alterations in the recurrence had a high degree of concordance to the original mass (Figure 6A), consistent with prior genomic work in this field (33–38). We further queried the CAF population in this patient, which was present in all three time points (Figure 6B). We observed that the myCAF population was heavily reduced post-chemotherapy, with lower expression in the recurrence as well (Figure 6C). However, both the iCAF and neural-CAF signatures were increased in the primary tumor site post-treatment, and in the recurrence timepoint, we saw predominant expression of the neural-CAF signature (Figure 6C). Comparison of the treatment-naïve epithelial cells to the recurrence epithelial cells revealed that genes involved in axonal guidance, Wnt signaling, and PI3K-Akt signaling were enriched in recurrent disease as compared to the treatment naïve primary tumor (Supplementary Figure 6A). This included axonal guidance genes such as *NRG3, NTN1, SEMA4G, and NRP1*, as well as receptors for non-canonical Wnt5a (*ROR1, RYK, LRP6*) (Supplementary Figure 6A). This shift towards a neural-like phenotype in cancer recurrence that mirrors the enrichment of neural genes and non-canonical Wnt mediators in neural-CAFs could be important in the context of perineural invasion and is consistent with previous reports of neural-like malignant cells found in PDAC (7,39).

**Figure 6.**
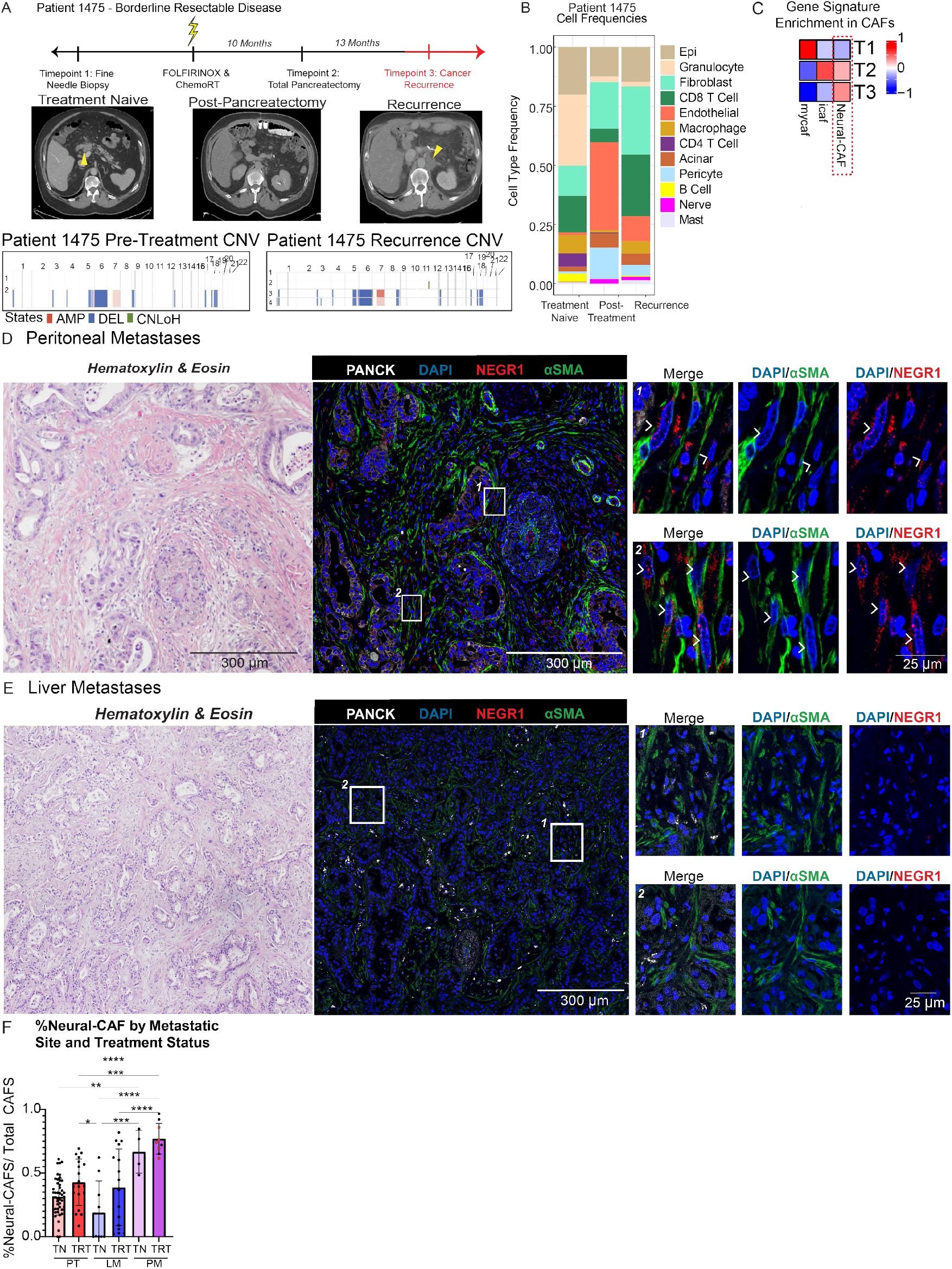
PDAC recurrence displays synchronous enrichment of neural genes in CAFs, and local metastases. **A) Top:** Timeline showing patient 1475 fine-needle biopsy (FNB), neoadjuvant therapy, surgical resection, and identification of recurrent disease (top). **Center:** Computed Tomography scan of patient 1475 at time of diagnosis (left), post-surgery (middle), and at time of discovered cancer recurrence (right) - the tumor mass is denoted by yellow arrows. **Bottom**: Heatmap of inferred copy number variation analysis of treatment naïve FNB and recurrence. **B)** Bar plot comparing cell type frequencies of patient 1475 specifically split between treatment naïve FNB, surgical sample post-therapy at time of resection, and recurrence biopsy. **C)** Heatmap of average axonal-guidance gene signature expression split by treatment status in patient-1475 (T1 – treatment naïve, T2 – treated, T3 - recurrence). **D)** Hematoxylin and eosin (HE) staining of biopsy from peritoneal metastases of PDAC (left), and immunofluorescence staining of serial section from same patient tissue with DAPI, aSMA (green), PANCK (white), and NEGR1 (red)(middle). White arrows denote NEGR1+αSMA+ CAFs at higher magnification (right). **E)** HE staining of biopsy from liver metastases of PDAC (left), and immunofluorescence staining of serial section from same patient tissue with DAPI, aSMA (green), PANCK (white), and NEGR1 (red)(middle). **F)** Bar plot of percentage of neural-CAFs in treatment naïve (TN) primary tumor (PT), treated (TRT) PT, TN liver metastases (LM), TRT LM, and TN peritoneal metastases (PM), and TRT PM. (N=11 TN PT, N=6 TRT PT, N=2 TN LM, N=4 TRT LM, and N=3 TN PM, and N=9 TRT PM with 1-7 regions of interest selected for each). **G)** Adjusted P value for %neural-CAFs in: TN PT versus TN PM *< 0*.*01*, TN PT versus TRT PM *< 0*.*0001*, TRT PT versus TN LM *< 0*.*05*, TRT PT versus TRT LM *< 0*.*001*, TN LM versus TN PM *< 0*.*001*, TRT LM versus TN PM *< 0*.*001*, and TRT LM versus TRT PM *< 0*.*001*.

To further parse the potential for neural-CAFs to drive tumor progression, we stained patient derived tissues from biopsies of peritoneal metastases and liver metastases (Figure 6D, E, Supplemental Figure 6B, C) and observed high neural-CAF localization to peritoneal metastases specifically (Figure 6D, F, Supplementary Figure 6B) in comparison to primary tumors (both treated and untreated) (Figure 6F) and liver metastases (both treated and untreated) (Figure 6E, F, Supplementary Figure 6C). While peritoneal metastases are thought to occur through transcoelomic spread, as opposed to hematogenous spread via portal vein dissemination in liver metastases (40,41), the preferential enrichment of neural-CAFs in the former may suggest a specific role for perineural invasion in this metastatic process. Taken together, these data point to the potential role of neural-CAFs in promoting invasion and recurrence post-treatment through the crosstalk of axonal guidance and Wnt signaling pathways between nerves and tumor cells.

## Discussion

Here, in this study, we identified a sub-group of CAFs, termed neural-CAFs, that were characterized by high expression of neural axonal guidance genes, and found to be enriched post-treatment. While some neural-CAFs co-expressed myCAF/iCAF signatures (26), a large proportion represented a distinct population of CAFs. The neural-CAF population abundance was enriched in areas of perineural invasion regardless of treatment status but also increased in tumor distant to nerves in treated tissues. Coupled with the observation that neural gene expression in fibroblasts correlated with neural gene expression in tumor epithelial cells, these findings suggest the presence of a neural signaling axis between specific tumor cells and cancer-associated fibroblasts (CAFs) that can contribute to increased tumor cell survival.

PDAC has a uniquely high tropism for nerves, with perineural invasion and nerve hypertrophy being histopathological hallmarks of the disease (11). Even in the absence of disease, nerve hypertrophy and hypersensitivity have been identified in chronic pancreatic inflammation. Following PDAC initiation, cancerous cells have been identified in mouse models within sensory nerve ganglia and along the spinal cord due to this high level of neurotropism (42). Neural crosstalk has previously been established to mediate pancreatic pathogenesis in early stages of the disease, as evidenced by ablation of sensory neurons of the pancreas inhibiting PDAC tumorigenesis in mice (42–44). This is further supported by findings that axonal guidance signaling molecules play a role in PDAC growth and invasion, and that tumor cells “pseudo-synapse” to nerves due to these secreted factors and guidance cues (45).

We identified in our dataset that a portion of our neural-CAF gene signature was expressed in fibroblasts obtained from healthy donor pancreata, albeit at lower levels than chemotherapy-treated CAFs. CAFs have been identified previously to not only have various subtypes, but also to express differing neuronal genes indicating a high level of plasticity (7,46–48). This is especially true of neuron-associated fibroblasts, such as those within the epineurium that express high levels of axonal guidance genes, and which interact both with peripheral nerves and Schwan cells (49). The expression of only a portion of our neural-CAF signature genes indicate that while neuronal genes are recapitulated in healthy fibroblasts, likely due to populations responsible for nerve engagement, the acquisition of the full neural-CAF signature during malignant transformation may allow tumor cells to co-opt this pathway as a survival mechanism.

Neural-CAFs were highly expressed near perineural invasion regardless of treatment status but expanded into the overall TME following chemotherapy exposure. Given the prevalence of neural-ligand-receptor interactions between neural-CAFs and tumor cells, the neural-CAF expansion to all areas of tumor could be indicative of a therapy-induced polarization of the CAFs that is leveraged to provide tumor-promoting crosstalk. For example, axonal guidance molecules identified in our neural-CAF signature such as *NRP1*, the cognate co-receptor for semaphorins, have been reported in pancreatic cancer previously in the context of cell invasion, perineural invasion, and tumor growth (39,50–52). Semaphorin expression in tumor cells has been found to encourage differentiation of neuronal progenitors in the cancer stroma and establish the interface between nerves and cancer cells (11,39,53–56). Neuropilins have also been found to engage with *VEGF*β and *TGF*β, which are key growth factors involved in PDAC tumor progression (52). Other work has identified both neuronal gene-expressing CAFs and tumor cells to be enriched post-treatment utilizing unmatched patient biospecimens (7), as well as a neuroendocrine subtype of tumor cells in PDAC associated with chemoresistance and increased Myc expression (57). This finding was recapitulated in our data given increased *Myc* expression in our neural-CAF population. Neural-CAFs were also characterized by an enrichment in non-canonical Wnt signaling, a pathway that has been studied in the context of PDAC tumorigenesis, and found to drive both the cellular identity of precancerous lesions in PDAC, as well as a more aggressive cancer type that is resistant to the inhibition of KRAS (58). These studies, in conjunction with our findings, further highlight the potential for neuronal signaling in CAFs to drive resistance to cytotoxic therapies.

While our dataset has strength in its matched longitudinal samples, the dataset was limited in its small size (n=5), as well as the heterogeneity of treatment and disease progression across patients. Furthermore, through analysis of inferred copy number variations (12), we discovered a limited number of tumor epithelial cells present in our dataset post-treatment. We did, however, have the opportunity to study a subset of tumor epithelial cells due to the presence of a recurrence. We observed that the recurrent tumor cells highly expressed genes involved in axonal guidance, many of which have been reported in connection to biomarkers of perineural invasion in pancreatic cancer (39,59). The findings of axonal guidance genes in the tumor epithelial compartment such as *NTN1, NTN4, NRP1*, and *SEMA4G* was highly encouraging, given that the recurrence was local (as opposed to distant), and thus perineural invasion was the possible mechanism of tumor escape. Furthermore, the receptors for the non-canonical Wnt ligand Wnt5a (*ROR1, ROR2, RYK*) were highly expressed in the recurrence tumor epithelial cells, mirroring enrichment for non-canonical Wnt signaling found in neural-CAFs.

Given the small sample size of our cohort, we were able to expand our studies through the use of immunostaining of archival metastatic samples, including those obtained in recurrence. Interestingly, we found that local metastases to the peritoneal cavity had the highest percentage of neural-CAFs, further supporting the role of neural-CAFs in promoting cancer metastasis and recurrence. Whether their enrichment in peritoneal metastases reflects perineural invasion that enables transcoelomic spread or represents a specific polarization toward the neural-CAF subtype following metastatic establishment in the peritoneum, will be elucidated through future functional studies. Overall, the presence of both an increase in an axonal guidance signature in tumor cells and a consistent significant increase of increased neural-CAFs post-treatment indicates the importance of axonal guidance and neuronal genes in pancreatic cancer. Furthermore, the spatial localization of neural-CAFs near areas of perineural invasion and peritoneal metastases in pancreatic cancer patients leads us to hypothesize a role for these CAFs in cancer-nerve crosstalk, metastasis and invasion. The dynamic changes in neural-CAF prevalence in the TME over the course of PDAC progression opens up the question of whether treatment with chemotherapy may inadvertently lead to changes in the TME that can promote perineural invasion of surviving tumor epithelial cells and thus cancer recurrence, and if so, what treatment strategies can be utilized for the prevention of such outcomes.

## Methods

### Sample Collection

Human subjects’ research and sample collection was conducted in accordance with the ethical guidelines outlined in the Belmont Report with approval by the Institutional Review Board at the University of Michigan (Ann Arbor, MI; IRB HUM00025339) and at Mayo Clinic (Rochester, MN; IRB 66-06). Medical charts were reviewed to assess potential study patients who specifically had resectable, borderline resectable or locally advanced disease. These patients had the potential for surgery or fiducial marker placement, allowing for collection of a post-chemotherapy timepoint (T2). Tumor tissue was collected via surgical resection or fine needle biopsy as described previously (31,60). With respect to healthy donor pancreas tissues, acquisition of donor pancreata for research purposes was approved by the Gift of Life research review group.

### Single-Cell RNA Sequencing

Newly acquired samples were processed as described previously (14,31,60) and single-cell cDNA libraries were prepared and sequenced at the University of Michigan Advanced Genomics Core using the 10x Genomics Platform. Samples were run using 28×151 bp of sequencing to a depth of 100,000 reads according to the manufacturer’s protocol with either NovaSeq or NovaSeqXPlus (Illumina) to generate de-multiplexed Fastq files. Convert Conversion Software v3.9.3 or v4.0 (Illumina) was used to generate de-multiplexed Fastq files. All files were processed and realigned with Cell Ranger v7.1.0 and aligned to the GRC38 transcriptome. Pre-processing and analysis were done using R version v4.4.0, RStudio version v2025.5.999.999, and Seurat version v5.1.0.(61) Samples were merged, batch corrected, filtered, and normalized as previously done (14,31,60). Variable genes were identified via FindVariableFeatures() and then principle component analysis was applied via RunPCA() with the genes defined in the preceding step. Cell clusters were identified as described previously (14,31). Gene signatures scores (myCAF, iCAF, and immune markers of activation and exhaustion) were obtained from literature and gene set scoring was calculated on each cellular cluster using AUCell() as described (31,62). Copy number variations were inferred using Numbat (v1.4.2) and applied on all samples, this pipeline utilized both gene expression and germline heterozygous SNPs to generate the copy number variation probability for each cell (12). Clones were inspected for each patient sample and marked as normal or tumor prior. Data from healthy normal pancreas samples were used as a reference (31). We performed functional annotation using the Kyoto Encyclopedia of Genes and Genomes (KEGG) enrichment analysis applied through the clusterProfiler package (RRID:SCR_016884) and EnrichKegg(). Further analysis on the larger dataset was done by merging our generated data with human single-cell RNA sequencing (scRNA-seq) data that were previously published (31). Differential cell–cell communication was done using CellChat (17). Cell–cell communication was calculated between each cell-type pair using the manually curated interaction database (CellChatDB) on the merged dataset described above. Inferred ligand-receptor pairs were calculated for specific pathways of interest. Pseudobulk analysis was performed on each cell type as described previously (31).

### Immunofluorescence Assessment

Formalin-fixed, paraffin-embedded tissue sections were processed for immunofluorescence staining following rehydration and antibody incubation steps. Slides were baked at 60 °C for 2 hours, cooled to room temperature, and deparaffinized with two 5-minute washes in Histoclear (Cat# 64111-04). Slides were then rehydrated through two 5-minute washes in 100% ethanol, one 5-minute wash in 95% ethanol, and one 5-minute wash in 70% ethanol, followed by two 2-minute washes in running deionized water. Antigen retrieval was performed by immersing slides in preheated 1X citrate buffer (Cat# HK080-9K) and steamed for 20 minutes. Slides were cooled to room temperature on ice for 15 minutes, washed twice for two minutes in deionized water, and then 5 minutes in 1X Phosphate Buffered Saline (PBS). A hydrophobic barrier was drawn around the tissue sections using an A-PAP pen. Slides were incubated for 60 minutes at room temperature in blocking buffer (1% bovine serum albumin, 0.3 M glycine, 10.0% goat serum, 0.1% Tween 20 in 1X PBS).

Following blocking, slides were incubated overnight at 4 °C with 150 μL of the primary antibody diluted in blocking buffer. Primary antibodies used were NEGR1 (Abcam, Cat# ab96071, RRID: AB_10679337, Rabbit, 1:200), NRP1 (Abcam, Cat# ab16786, RRID: AB_2298830, Rabbit, 1:200), E-Cadherin (Cell Signaling Technology, Cat# 14472, RRID: AB_2728770, 1:300), Caspase 3 (Cell Signaling Technology, Cat# 9664, RRID: AB_2070042, Rabbit, 1:200) and MKI67 (Cell Signaling Technology, Cat# 12075, RRID: AB_2728830, Mouse, 1:200). The following day, slides were washed three times for five minutes each in 1X PBS and incubated for 1 hour at room temperature in the dark with fluorophore-conjugated secondary antibody diluted in blocking buffer. Secondaries used included Goat Anti-mouse AF488 (Cat# A-11001, 1:500, RRID:AB_2534069) and Goat Anti-Rabbit AF647 (Cat# A-21245, 1:500, RRID:AB_2535813).Slides were then washed three times for five minutes in 1X PBS and incubated overnight in fluorophore conjugated primary antibody (Pan-Cytokeratin (PanCK) Alexa Fluor 570 (Fisher Scientific, Cat# 50-112-3641, Mouse, pre-conjugated, 1:800) and α-Smooth Muscle Actin (αSMA) Alexa Fluor 555 (Abcam, Cat# ab184675, RRID: AB_2832195, Rabbit, pre-conjugated, 1:800)) diluted in blocking buffer. The following day slides were washed three times for five minutes each in 1X PBS and counterstained with DAPI (1:10000 in 1X PBS, for 5 minutes at room temperature. After three additional PBS washes, slides were mounted with ProLong™ Diamond Antifade (Cat# P36934), cover slipped and allowed to dry overnight in the dark. Slides were stored at 4 °C until imaging. Slides were imaged on Nikon Ti2 Widefield microscope at 10X and Leica Stellaris Confocal 5 at 40X with oil.

### Immunofluorescence Assessment with Tyramide Signal Amplification

Slides were processed the same as immunofluorescent assessment (detailed above) but with modifications per Alexa Fluor 488 Tyramide SuperBoost Kit instructions (Cat# B40922). After antigen retrieval, slides were cooled to room temperature on ice for 15 minutes, washed twice for two minutes in deionized water. Slides were incubated for 15 minutes in humidified chamber with 3 drops of kit provided 3% hydrogen peroxide and then washed in 1X Phosphate Buffered Saline (PBS). Slides were then blocked for 30 minutes at room temperature with provided blocking buffer (Component A). A hydrophobic barrier was drawn around the tissue sections using an A-PAP pen. Primary antibody for signal amplification was diluted in 1X PBS (Anti-PDGFR alpha + PDGFR beta antibody [Y92] – C terminal, Abcam, CAT#: ab32570, Rabbit, 1:500, RRID:AB_777165) and incubated in 4-degrees C overnight. The next day, slides were washed in 1X PBS and 2-3 drops of provided Poly-HRP conjugated secondary antibody was added for 60 minutes at room temperature. Slides were washed in 1X PBS. Tyramide working solution was prepared from provided reagents in kit (100X Tyramide stock solution (1:100), 100x H202 solution (1:100), 1X reaction buffer). 100 μl of the working solution was incubated on samples for 8 minutes in the dark. 100 μl of the provided working stop reagent (1:11 dilution in PBS) was added directly to slides and incubated for 5 minutes in the dark. Slides were washed in 1X PBS. Immunofluorescence assessment protocol was continued from this step starting at antigen retrieval step (described above) for remainder of antibodies.

### Image Analysis and Quantification

Images were quantified using FIJI (v2.14.01) and Python scripts run in Jupyter Notebook (v3.13.7). Full 4-channel.*ome*.*tif* files were pulled into FIJI and split into single channel images. Regions of interest were chosen using pathologist marked hematoxylin and eosin slides and then copied over to the DAPI channel for each fluorescence image. These ROIs were saved in FIJI, and a macro was used to apply settings to all channels and multi crop into 10 single channel regions of interest per tissue. These cropped files were read into a python script that allowed users to identify structures of interest to be analyzed. These polygons were then imported, along with all cropped single channel files into Jupyter Notebook and each channel was masked individually and then double positive masks were created for all CAFs (DAPI+αSMA+), neural CAFs (DAPI+αSMA+NEGR1+ and DAPI+αSMA+NRP1+), and proliferating tumor cells (DAPI+PANCK+KI67+) and all tumor cells (DAPI+PANCK+). Percent neural CAFs and percent proliferating tumor cells were calculated as a percentage of total CAFs and total tumor cells. ROIs were marked as having tumor-nerve involvement – identified as areas where tumor cells and nerves were within 300 μm of each other, or having just nerves in healthy stromal tissue, or tumor cells distant from nerve. Percentages of each cell type were graphed with all ROIs merged or split by structure and/or treatment status of the patient. We stained n=11 treatment naïve and n=10 treated patient tissues, but due to poor tissue quality of some patients we quantified n=11 treatment naïve and n=6 treated patients. In the python script, double positive cells were calculated in outward radiuses starting from the border of the identified structure (either a nerve, or a path of tumor cells) at 50 μm distances up to 300 μm away from the structure border. For overall differences in structures and treatment, all radial bins were summed. Neural CAFs were also calculated as a function of distance from the structure and graphed in this way. Significance was calculated in GraphPad Prism 10 with a two-tailed student t-test between categories of interest. Proliferating tumor cells were graphed on a patient specific basic between areas of perineural invasion and tumor cells (distant to nerve) and analyzed with a paired t-test. For metastatic biopsies, large tissues of tumor were broken down into 4 randomly selected regions of interest, and small biopsies were taken as one region of interest in their entirety. These were processed and quantified in the same manner as the treated and treatment naïve primary tumor tissues. Significance was calculated in GraphPad Prism 10 with a one-way ANOVA and multiple comparisons between the mean of each sample group with the mean of every other sample group.

## Data Availability

Raw human data from the Carpenter, et. al. Study[GOL reference] are available NIH Gene Expression Omnibus database GSE229413. Additional unpublished sequencing files in this manuscript will be uploaded to new study accessions upon acceptance.

## Code Availability

All code used for this study will be made publicly available at the following GitHub upon acceptance.

## Statistical Analysis

Statistical analysis were either performed with RStudio for scRNA-seq analysis utilizing a Wilcoxon rank sum test from the ggpubr package with stat_compare_means() or in GraphPad Prism 10 with a two-tailed student t-test between groups of interest, a paired t-test for within patient comparisons, and a one-way ANOVA for multiple comparisons.

## Acknowledgements

We sincerely thank all patients and organ donors who have generously provided tissue for this research. We thank D. Postiff from the University of Michigan Tissue Procurement Core for procurement of surgical samples. We thank T. Tamsen, M. Hogan and J. Opp from the University of Michigan Advanced Genomics Core for their assistance with single cell RNA sequencing. We thank the University of Michigan Advanced Microscopy Core for assistance acquiring immunofluorescent images. We thank Hamadi Madhi for his assistance with single cell sequencing library preparation. We thank Christopher Strayhorn for histology services. We acknowledge the Rogel and Blondy Center for Pancreatic Cancer for its support of this study. Work in Carpenter Lab was supported by the Department of Veterans Affairs BLRD Career Development Award CDA IK2BX005875, the American College of Gastroenterology Career Development Award, the Research Scouts Award by the University of Michigan, and the Cancer Discovery Award by the University of Michigan Rogel Cancer Center. A.Z.H was supported by NIGMS GM150581. P.K. was supported by NIH/NIAID T32-AI007413 and NIH/NIGMS T32-GM113900. J.S was supported by R37CA262209. The funders had no role in study design, data collection and analysis, decision to publish or preparation of the manuscript.

