## supplemental figures and legends for "Longitudinal Analysis of Matched Patient Biospecimens Reveals Neural Reprogramming of Cancer-Associated Fibroblasts Following Chemotherapy in Pancreatic Cancer"

Supplementary Figure 1

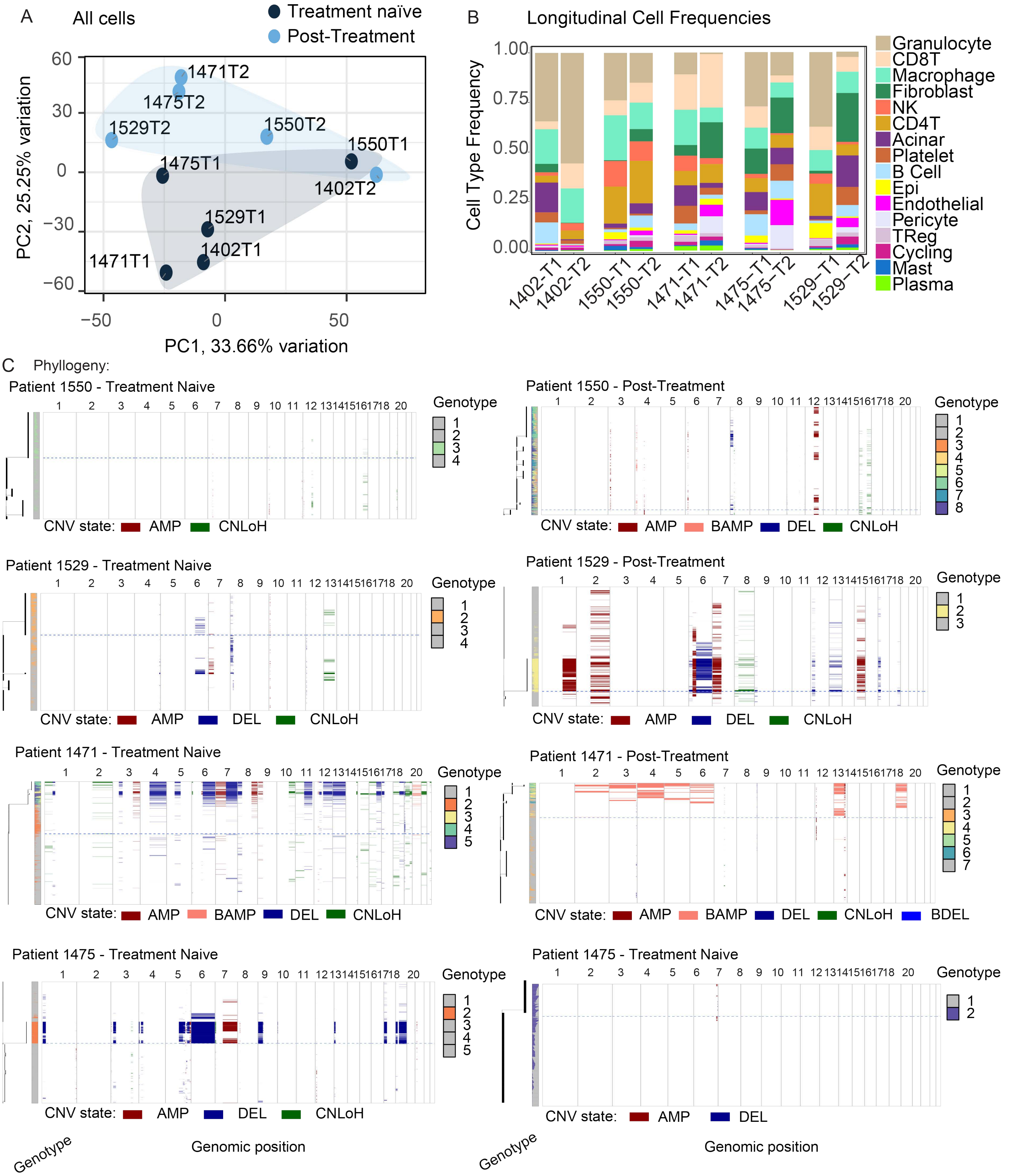

Supplementary Figure 1: Longitudinal profiling of PDAC tumors reveals global transcriptomic differences with therapy.

- **A)** PCA plot of pseudobulk-aggregated counts from each sample. Each dot represents one aggregated single-cell sequencing sample.
- **B)** Histogram of cell-type abundance of all cell populations by patient and treatment status.
- **C)** Single-cell CNV landscape and reconstructed phylogeny of four patients (1550, 1529, 1471, 1475) before (left) and after (right) chemotherapy treatment. The legend included indicates the identified variant as an amplification (AMP), balanced amplification (BAMP), deletion (DEL), balanced deletion (BDEL), or CNLoH (copy-neutral loss of heterozygosity).

Supplementary Figure 2

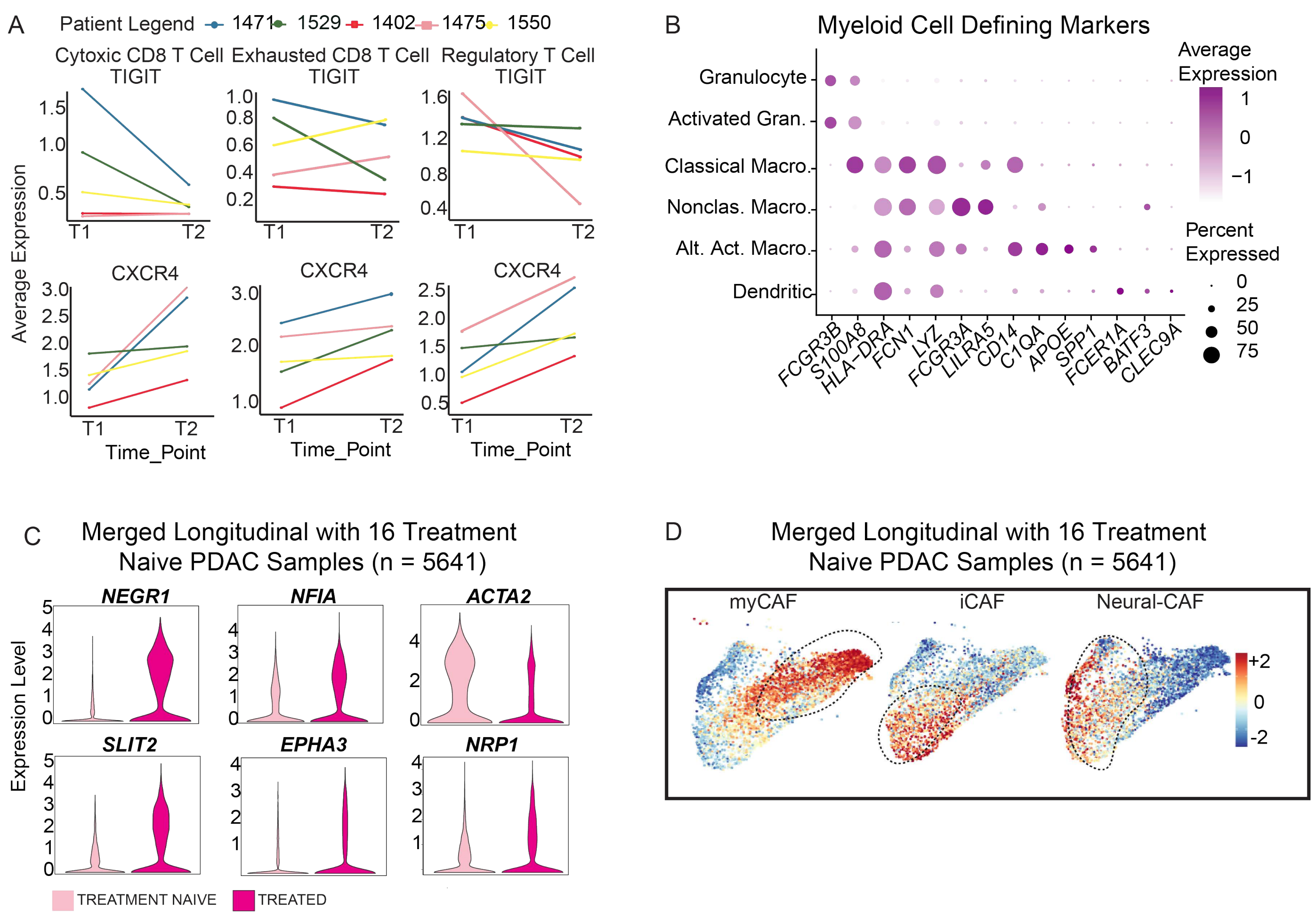

Supplementary Figure 2: Longitudinal profiling of PDAC tumors reveals global transcriptomic differences with therapy in immune cell component and an ‘axonal guidance’ gene signature is enriched in a sub-population of CAFs following treatment in a larger dataset.

- **A)** Before and after plots of *TIGIT* and *CXCR4* in T cell populations.
- **B)** Dot plot of identifying genes in each myeloid cell population.
- **C)** Violin plots of neural-CAF genes and *ACTA2* in treatment naïve PDAC samples (T1) and treated PDAC samples (T2) in the merged dataset. Adjusted *P* value for significantly differentially expressed markers: *NRP1 <2.2e-16, NEGR1 <2.2e-16, NFIA <2.2e-16, SLIT2 <2.2e-16, EPHA3 3.2e-11, ACTA2 <2.2e16.*
- **D)** Average gene expression of myofibroblastic genes (left), inflammatory fibroblast genes (middle), and neural-CAF genes (right) mapped to fibroblasts in merged dataset.

Supplementary Figure 3

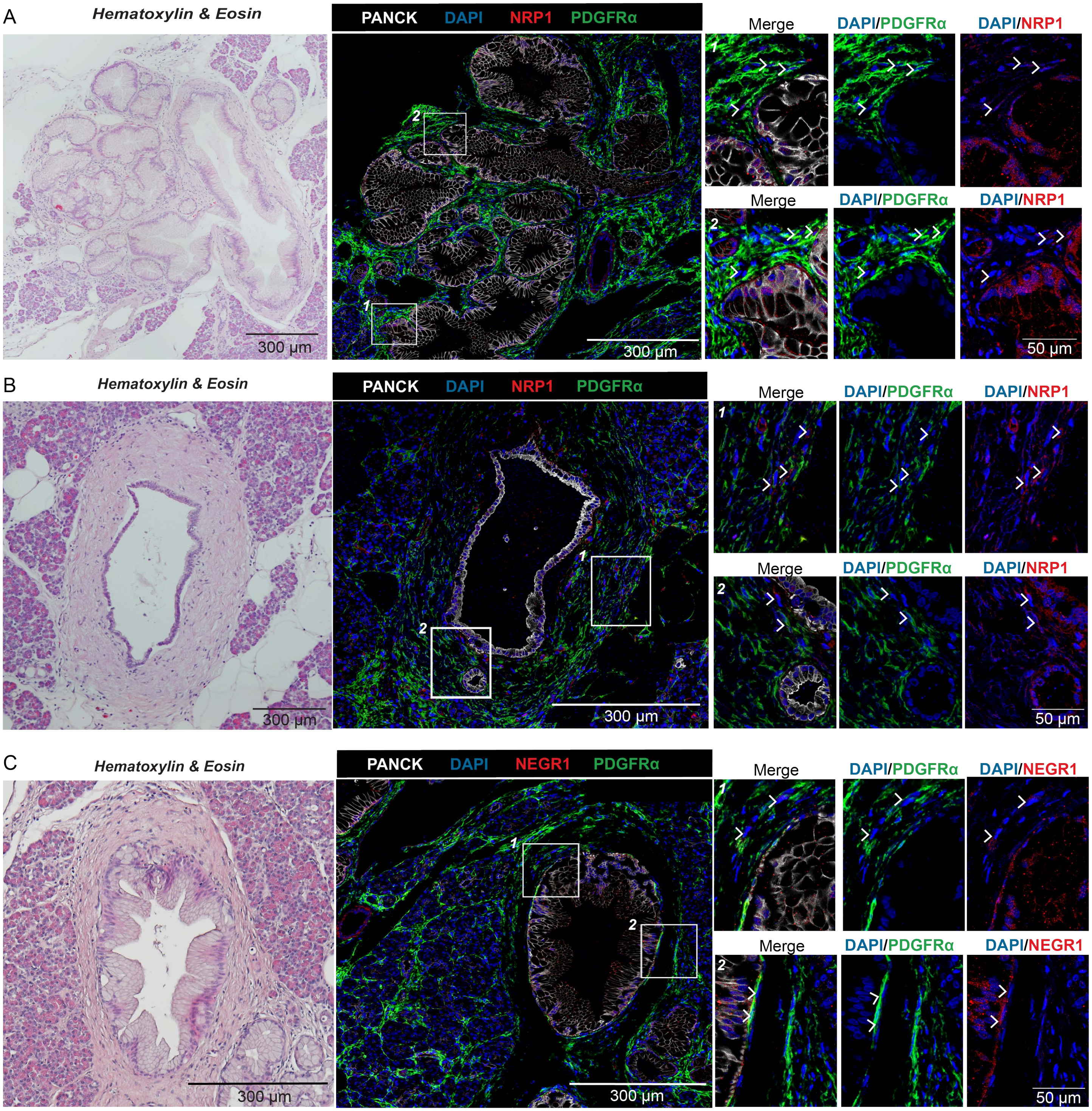

Supplementary Figure 3: Neural-CAF population in partially recapitulated in healthy pancreata.

**A)** (Left) Hematoxylin and eosin (HE) staining of human donor tissue. (Right) Immunofluorescence staining of serial section with DAPI, PDGFRα (green), pan-CK (white), and NEGR1 (red) in an area identified as a PanIN. White arrows denote NEGR1+PDGFRα+ fibroblasts.

**B)** (Left) HE staining of human donor tissue. (Right) Immunofluorescence staining of serial section with DAPI, PDGFRα (green), pan-CK (white), and NRP1 (red) in an area identified as a normal duct. White arrows denote NRP1+PDGFRα+ fibroblasts.

**C)** (Left) HE staining of human donor tissue. (Right) Immunofluorescence staining of serial section from patient with DAPI, PDGFRα (green), pan-CK (white), and NEGR1 (red) at 40X in an area identified as a normal duct. White arrows denote NEGR1+PDGFRα+ fibroblasts.

Supplementary Figure 4

A Patient 34800 (Treatment Naive)

Perineural Invasion | Tumor (distant from nerve) | Nerve (distant from tumor)

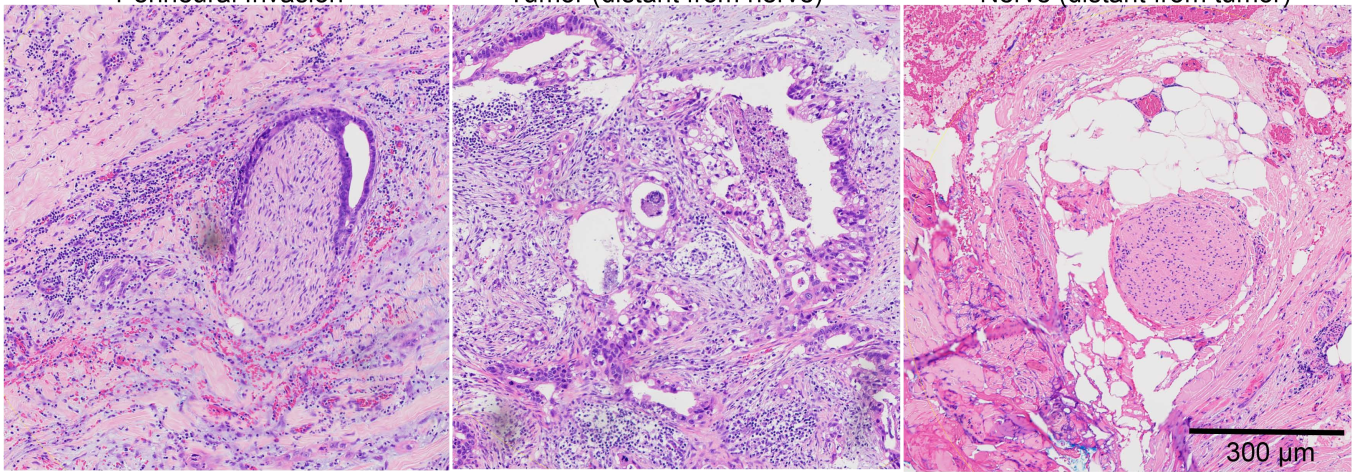


B Patient 51536 (Treated)

Perineural Invasion | Tumor (distant from nerve) | Nerve (distant from tumor)

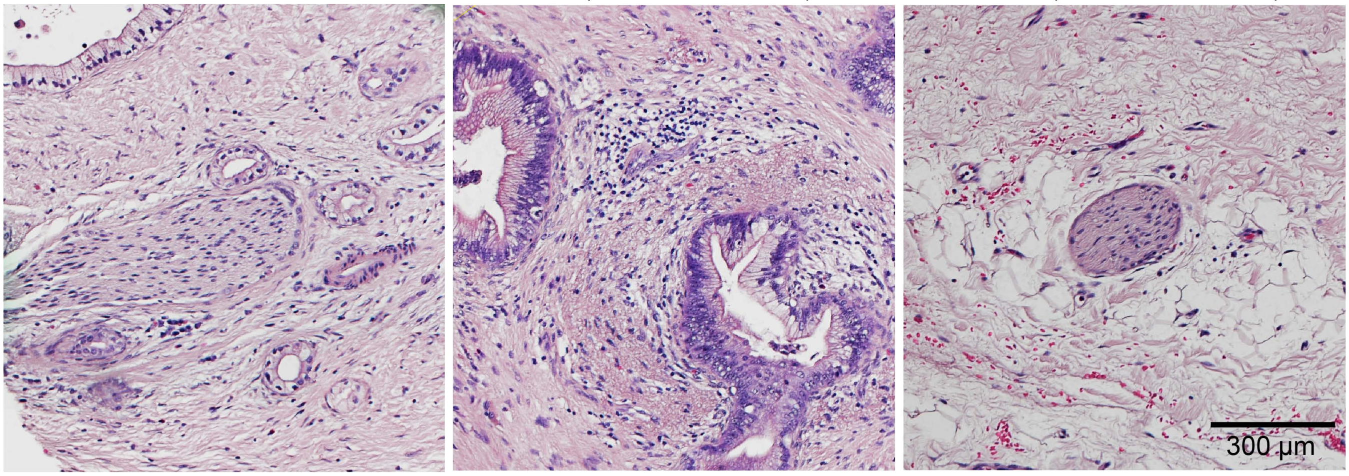


C Radial Bins for Quantification (Pt. 1926)

**Perineural Invasion** | **Tumor (distant to nerve)**

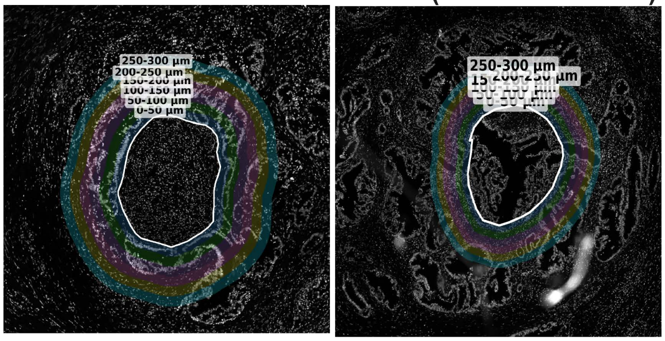


D KI67 expression near PNI v. Tumor Only in Pt. 1926

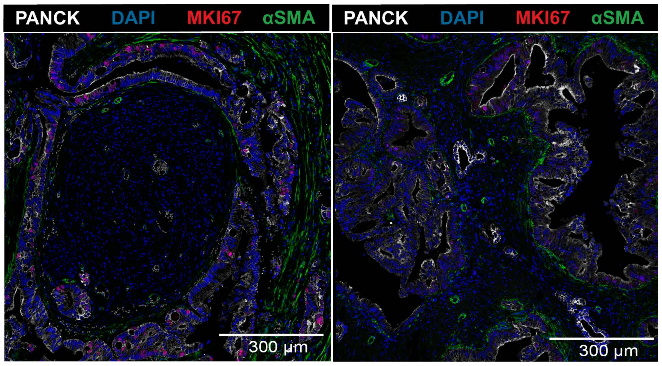

Supplementary Figure 4: Regions of perineural invasion, tumor (distant from nerve) and nerve (distant from tumor) were identified in treatment naïve and treated patients and quantified for percent neural-CAFs.

- **A)** (Left) Hematoxylin and eosin (HE) staining of treatment naïve PDAC patient tissue (34800) in an area of perineural invasion (PNI) (Center). HE staining of treatment naïve PDAC patient tissue (34800) in an area of tumor (distant from nerve) (Right). HE staining of treatment naïve PDAC patient tissue (34800) in an area of nerve (distant from tumor).
- **B)** (Left) HE staining of treated PDAC patient tissue (51536) in an area of PNI (Center). HE staining of treated PDAC patient tissue (51536) in an area of tumor (distant from nerve) (Right). HE staining of treated PDAC patient tissue (51536) in an area of nerve (distant from tumor).
- **C)** Immunofluorescent images of DAPI from PNI region (left) of interest and tumor (distant to nerve) region (right) of interest. Identified polygon structure and radial bins surrounding each structure are shown. Bins are at 50-micron intervals from 0-300 microns away from the edge of the structure.
- **D)** Co-immunofluorescence of area of perineural invasion and tumor nonadjacent to perineural invasion from patient 1926 tissue stained for DAPI (blue), a-SMA (green), PANCK (white), and MKI67 (red)

Supplementary Figure 5

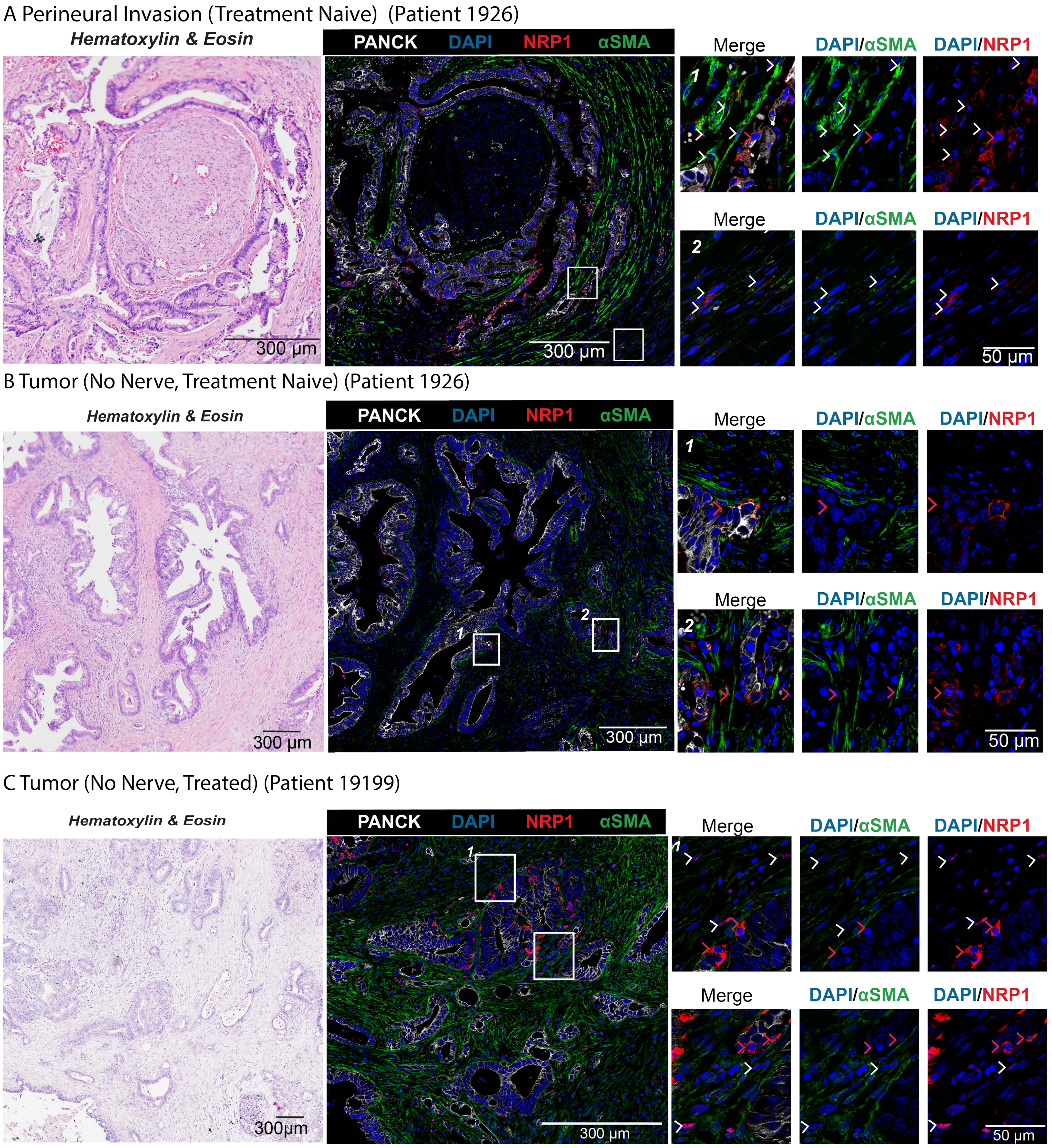

Supplementary Figure 5: NRP1+ CAFS are enriched near perineural invasion and in treated tumors.

**A)** (Left) Hematoxylin and eosin (HE) staining of treatment naïve PDAC patient tissue (1926). (Right). Immunofluorescence staining of serial section with DAPI, αSMA (green), pan-CK (white), and NRP1 (red) in an area marked as perineural invasion (PNI). White arrows denote NRP1+ CAFs and red arrows denote NRP1+ tumor cells in areas of higher magnification.

**B)** (Left) HE staining of treatment naïve PDAC patient tissue (1926) (Right). Immunofluorescence staining of serial with DAPI, αSMA (green), pan-CK (white), and NRP1 (red) at 40X in an area marked as tumor with no adjacent nerve. No NRP1+ CAFs were identified here, red arrows denote NRP1+ tumor cells in areas of higher magnification.

**C)** (Left) HE staining of treated PDAC patient tissue (19199) (Right). Immunofluorescence staining of serial section with DAPI, αSMA (green), pan-CK (white), and NRP1 (red) in an area marked as tumor with no adjacent nerve. White arrows denote NRP1+ CAFs and red arrows denote NRP1+ tumor cells in areas of higher magnification.

Supplementary Figure 6

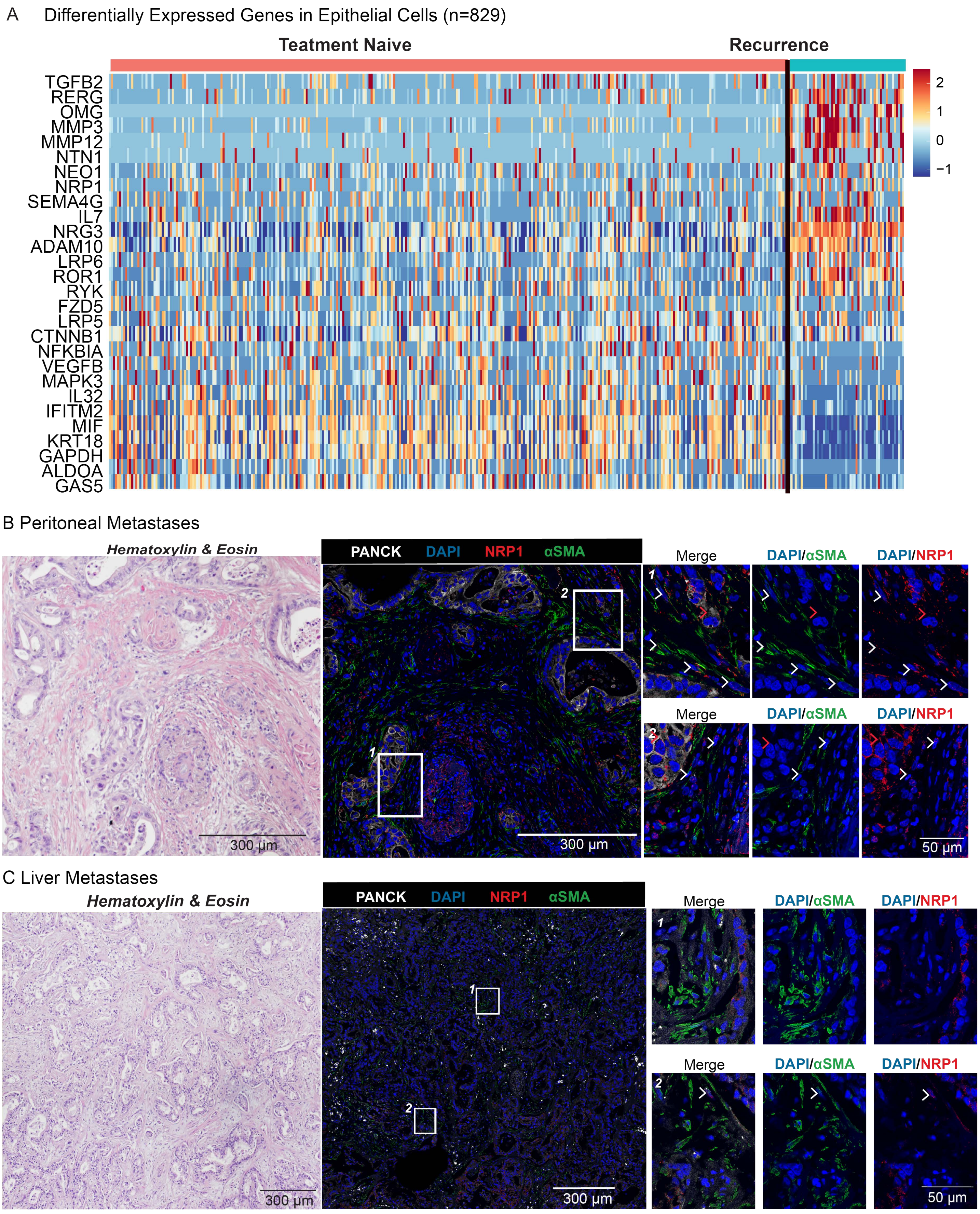

Supplementary Figure 6: PDAC recurrence displays synchronous enrichment of neural genes in tumor cells compared to the treatment naïve primary tumor.

- **A)** Top differentially expressed genes between tumor epithelial cells of patient 1475 treatment naïve FNB (left) and patient 1475 recurrence biopsy (right).
- **B)** (Left) Hematoxylin and eosin (HE) staining of patient biospecimen from a peritoneal metastases of PDAC patient tissue (Right). Immunofluorescence staining of serial section with DAPI, αSMA (green), pan-CK (white), and NRP1 (red). White arrows denote NRP1+ CAFs in areas of higher magnification.
- **C)** (Left) HE staining of patient biospecimen from a liver metastases of PDAC patient tissue. (Right). Immunofluorescence staining of serial section with DAPI, αSMA (green), pan-CK (white), and NRP1 (red). White arrows denote NRP1+ CAFs in areas of higher magnification.
